# Mechanical Skin Stress-Induced Lesion Development via ATP-Amplified Neutrophil Extracellular Trap Formation

**DOI:** 10.64898/2026.03.11.710999

**Authors:** Daisuke Matsumoto, Katja Bieber, Ralf J. Ludwig, Daisuke Tsuruta, Sho Hiroyasu

**Affiliations:** Department of Dermatology, Graduate School of Medicine, Osaka Metropolitan University, 1-4-3 Asahi-machi, Abeno-ku, 545-8585, Osaka, Japan; Lübeck Institute of Experimental Dermatology, University of Lübeck, Ratzeburger Allee 160/Haus B9, 23562, Lübeck, Germany

**Keywords:** Neutrophils, Neutrophil extracellular traps, ATP, Complement components, Mechanical skin stress, Neutrophilic skin diseases, Epidermolysis bullosa acquisita

## Abstract

Neutrophilic skin diseases, including Behçet disease, Sweet syndrome, pyoderma gangrenosum (PG), and epidermolysis bullosa acquisita (EBA), are characterized by an exaggerated inflammatory response following mechanical skin stimulation, yet the underlying mechanisms remain unclear. We identify adenosine triphosphate (ATP) released from keratinocytes as a key mediator of this phenomenon, promoting neutrophil extracellular trap (NET) formation. Using an EBA murine model as a model of neutrophilic skin disease, where scratching (a prototypic mechanical stimulation) exacerbates lesional severity, we observed abundant NET deposition in lesional skin. Degradation of these NETs with DNase1 reduced clinical and histopathological severities. In vitro, purified NET components increased IL-8 secretion from keratinocytes and fibroblasts, suggesting that NETs amplify inflammation via a self-amplifying loop of neutrophil recruitment. In the EBA mouse, scratch restriction with neck collars not only attenuated clinical and histological disease severities but also decreased lesional NETosis and neutrophils. Mechanistically, keratinocytes released ATP in response to mechanical stress in vitro, and pharmacologic purinergic blockade in the EBA mice with suramin phenocopied the protective effects of scratch restriction. While ATP alone did not induce NETosis, ATP enhanced complement component 5a (C5a)-induced NET formation in vitro. These findings indicate that keratinocyte-derived ATP, released in response to mechanical stress, contributes to NETosis in a C5a-dependent manner, thereby exaggerating neutrophilic inflammation, leading to blistering and further NETosis. Histopathological analyses of EBA and PG cases also demonstrated NETs accumulation localized to the upper dermis, suggesting a conserved ATP-NET axis. Targeting this pathway may represent a promising therapeutic strategy for neutrophilic skin diseases.

## 1. Introduction

Neutrophilic skin diseases, characterized by prominent neutrophilic infiltration in the lesional dermis, encompass several intractable conditions such as Behçet disease, Sweet syndrome, pyoderma gangrenosum (PG), and epidermolysis bullosa acquisita (EBA) [1, 2]. These disorders share clinical characteristics including resistance to conventional therapies, responsiveness to specific reagents such as dapsone and colchicine, and a hypersensitivity to mechanical stimulation, known as pathergy [1, 3, 4]. Due to the neutrophilic infiltration in lesional skin, induction of neutrophil extracellular traps (NETosis) has been suspected in these diseases, however, direct evidence remains limited [5].

Neutrophils release neutrophil extracellular traps (NETs), web-like structures composed of citrullinated histones and granule-derived proteins, in response to a variety of stimulations including nicotinamide adenine dinucleotide phosphate (NADPH) oxidase activation, toll-like receptor signaling, and complement activation [6–9]. During this process, termed NETosis, protein arginine deiminase 4 (PAD4) catalyzes the citrullination of arginine residues in the core histones (H2A, H2B, H3, and H4), leading to chromatin decondensation and the subsequent release of nuclear deoxyribonucleic acid (DNA) along with histones and various granule-derived proteins into extracellular spaces [10, 11]. NETs play a critical role in host defense by trapping microorganisms and by modulating immune responses during infection. Moreover, accumulating evidence implicates NETs in the pathogenesis of various inflammatory disorders, including psoriasis, systemic lupus erythematosus, rheumatoid arthritis, and Stevens-Johnson syndrome/toxic epidermal necrolysis, through mechanisms such as inflammatory cytokine production, T cell regulation, and B cell activation [8, 12–15]. However, the precise role of NETs in neutrophilic skin conditions remains incompletely understood, partly due to the lack of suitable animal models that recapitulate the key features of human neutrophilic skin diseases.

EBA is a prototypical neutrophilic skin disease characterized by prominent neutrophilic inflammation and subepidermal blistering [2, 16]. Its pathogenesis is primarily induced by autoantibodies against collagen VII (COL7), a key molecule of the dermal-epidermal attachment, followed by complement activation and neutrophilic infiltration. Passive transfer of rabbit IgG against mouse Col7 induces an EBA-like phenotype in mice, featuring robust neutrophilic infiltration and subepidermal blistering [17]. This response is complement-dependent, with a dominant contribution from the alternative pathway [18]. Although NETs have been detected in pemphigoid diseases (the broader disease category that includes EBA) [15, 19], the pathological role of NETs in these conditions, including EBA specifically, remains unclear. Importantly, the passive IgG transfer EBA model captures key clinical and immunological features of neutrophilic skin diseases, such as prominent neutrophilic infiltration and pathergy-like exacerbation by scratching, while providing quantifiable clinical and histopathological severity metrics, making it an experimentally tractable system to test NET-centered mechanisms in neutrophilic skin diseases.

In this study, using the passive IgG transfer EBA murine model, we demonstrate that NETosis occurs in lesional skin and that its inhibition ameliorates disease severity. Mechanistically, purified NET components induced IL-8 secretion from both keratinocytes and fibroblasts, likely contributing to sustained neutrophilic inflammation. Adenosine triphosphate (ATP) released from physically stressed keratinocytes, in combination with activated complement stimulation, triggers NETosis both in vivo and in vitro. Furthermore, in human patients, NETosis was found to be predominantly localized near the epidermis not only in EBA but also in PG, suggesting a conserved pathogenic mechanism in neutrophilic skin diseases. Taken together, these findings elucidate a previously unrecognized pathway in which mechanical stimulation induces keratinocyte-derived ATP release, leading to NETosis and persistent neutrophilic infiltration, suggesting a shared mechanism across neutrophilic skin diseases.

## 2. Materials and Methods

### 2.1. Human samples

Peripheral blood samples from healthy controls were obtained from residual specimens remaining after routine clinical testing at Osaka Metropolitan University Hospital (Osaka, Japan). Written informed patient consent was obtained from all donors. Formalin-fixed paraffin-embedded lesional skin specimens were obtained from patients diagnosed with EBA, Behçet disease, Sweet syndrome, or PG based on laboratory, histopathological, and clinical findings. As these samples were collected prior to this study, the Institutional Review Board waived the requirement for written informed consent and permitted an opt-out process in accordance with institutional and international guidelines. All experimental procedures using human samples were approved by Osaka Metropolitan University Hospital Institutional Review Board (Approval No. 2024-165; January 29, 2025).

### 2.2. Laboratory mice

6-week-old female C57BL/6NCrSLC mice were obtained from Japan SLC Inc, (Shizuoka, Japan). Mice were housed in a 12-hour light/dark cycle with controlled room temperature at 22-26 °C and 25-40% relative humidity at a specific pathogen-free barrier facility in the Graduate School of Medicine at Osaka Metropolitan University. All experimental procedures using animals were approved by Osaka Metropolitan University Animal Experiment Committee.

### 2.3. Animal models

Systemic antibody-transfer murine model of EBA was established as described elsewhere [17]. Briefly, 7-week-old female mice received intraperitoneal injections of 150 μg rabbit anti-mouse Col7 IgG once every second day over 12 days. All mice were single-caged throughout the protocol. For the evaluation of lesional area in EBA mice, visible erythema, crusts, and ulcers were defined as lesions. The whole lesional area or facial lesional area were divided by the estimated full body surface area (9.82 × body weight (g) ^ 0.667) to quantify the percentage of lesional area, defined as the affected surface area, adapted from former literature [20]. The mice were euthanized on day 12 and skin samples were collected for analyses. For scratch prevention experiment, light and flexible polypropylene collars were placed around the neck of the mice to prevent scratching with their legs. The collars were applied 3 days prior to the administration of anti-Col7 IgG and placed until the study endpoint. In purinergic inhibition experiment, 100 μL of 4 mg/mL suramin sodium salt (S2671-100MG; Sigma-Aldrich, Inc., St. Louis, MO, U.S.) or 100 μL of 0.9% NaCl solution (K1F91; Otsuka Pharmaceutical Co, Ltd., Tokyo, Japan) were intraperitoneally administered daily for 12 consecutive days, starting one day before anti-Col7 IgG administration. In the NETosis inhibition experiment, 20 μL of 500 U/mL DNase1 (18068-015; Thermo Fisher Scientific, Inc., Waltham, Massachusetts, U.S.) or 50 μL of 0.9% NaCl solution were intraperitoneally administered daily for 12 consecutive days, starting one day before anti-Col7 IgG administration.

Sample sizes for all animal studies were determined by statistical power calculations prior to the experiments based on standard deviations estimated from our former or preliminary results.

### 2.4. Histological and immunohistochemical analyses

Five-μm sections of paraffin-embedded skin tissue blocks were deparaffinized and rehydrated for hematoxylin and eosin (H&E) and immunohistochemical analyses. Histological blister score was calculated from H&E-stained samples using the formula ((combined total length of all blistered regions)/(combined total length of all dermal-epidermal junction examined)) × 100, as adapted from a previously described method [21].

For immunohistochemistry (IHC), the sections were processed for antigen retrieval by boiling in target retrieval solution (S1699; Agilent Technologies, Inc., Santa Clara, California, U.S.), followed by blocking with 5% donkey serum (D9663; Sigma-Aldrich, Inc.) for 1 hour at room temperature. The sections were incubated overnight at 4 °C with primary antibodies; 1:200 dilution of anti-myeloperoxidase (MPO) antibody (AF3667; Bio-Techne Corporation., Minneapolis, Minnesota, U.S.) and 1:50 dilution for anti-citrulinated histone H3 (CitH) (citrulline R2 + R8 + R17) antibody (ab5103; Abcam., Cambridge Biomedical Campus, Cambridge, U.K.). The sections were subsequently incubated with Alexa Fluor 488- and 594-conjugated secondary antibodies (R37118, A11058; Thermo Fisher Scientific, Inc.) at a 1:500 dilution followed by the 5-minute incubation with 4, 6-diamidino-2-phenylindole (DAPI) (D1306; Thermo Fisher Scientific, Inc.).

Fluorescence images were acquired using the All-in-One Fluorescence Microscope (BZ-8000; Keyence, Osaka, Japan). Images of each section were analyzed using ImageJ (National Institutes of Health., Bethesda, Maryland, U.S.). For the evaluation of neutrophils and NETosis, the threshold function in ImageJ was applied, while carefully excluding the epidermis, hair follicles, and vesicles. NET area score ((NETosis area of upper or lower dermis)/(total observation area) × 100), neutrophil score ((neutrophil number)/(total observation area)), and NET index score ((NETosis area)/(neutrophil number)) were calculated from the obtained images. For human skin samples, quantification was applied separately for the upper dermis area (<400 μm from the dermal-epidermal junction) and the lower dermis area (>400 μm from the dermal-epidermal junction).

### 2.5. Cell culture

Human adult low calcium temperature keratinocytes (HaCaT) (300493F; Cell Lines Service GmbH., Eppelheim, Germany) and human fibroblast cells (PCS-201-030; American Type Culture Collection., Manassas, Virginia, U.S.) were cultured in Dulbecco’s Modified Eagle Medium (DMEM) (041-29775, 044-29765; FUJIFILM Wako Pure Chemical Corporation., Osaka, Japan) supplemented with 10% fetal bovine serum (10270-106; Thermo Fisher Scientific, Inc.), 100 U/mL penicillin and 100 μg/mL streptomycin (168-23191; FUJIFILM Wako Pure Chemical Corporation.). Cells were maintained at 37 °C in a 5% CO2 atmosphere and passaged using 0.05% trypsin-EDTA (202-16931; FUJIFILM Wako Pure Chemical Corporation.). Cultured cells were used for the NET component stimulation assay and scratch assay.

### 2.6. NET component and ATP stimulation assays

Neutrophils and serum were isolated from peripheral blood of healthy donors for use in NET component stimulation and ATP stimulation assays. To isolate the neutrophils, whole blood was layered in a 1:1 volume ratio on top of Polymorphprep (1114683; Abbott Laboratories., Abbott Park, Illinois, U.S.) and centrifuged without brake, and neutrophil layer were collected and washed according to the manufacturer’s instructions. Residual erythrocytes in the neutrophil layer were lysed by incubation with 10-fold-diluted Lysis buffer (60-00051-10; PluriSelect USA., El Cajon, California, U.S.) for 7 minutes. Neutrophils were washed and plated in 6 cm dishes at 1.0 × 10^6^ /mL in serum-free Roswell Park Memorial Institute 1640 (RPMI1640) medium (11875093; Thermo Fisher Scientific, Inc.).

For the cell-free NET component stimulation assay, 1.0 × 10^6^ cells/mL of the neutrophils were stimulated with 600 nM phorbol 12-myristate 13-acetate (PMA) (S7791; Selleck Biotechnology., Yokohama, Japan) for 4 hours at 37 °C. Following the stimulation, the medium was discarded to remove residual PMA, cytokines, and other soluble components. The cell layer was resuspended in phosphate-buffered saline (PBS) and centrifuged, and the resulting supernatants containing cell-free NET components were collected. As a control, supernatants from neutrophils incubated with vehicle for 4 hours were prepared in parallel. To assess the stimulatory effects of NET components, 0.02, 0.2, or 0.5 mL of NET component-containing or control supernatant was added to 5.0 × 10^5^ HaCaT cells or fibroblasts cultured in 12-well plates. After 12 hours of stimulation, the cultured supernatants were collected for quantification of IL-8 levels using a Human IL-8 enzyme-linked immunosorbent assay (ELISA) kit (ab214030; Abcam) according to the manufacturer’s protocol.

For the ATP stimulation assay, 100 μL of neutrophil suspension (5.0 × 10^5^ cells/mL) was added to glass-bottomed dish. Neutrophils were stimulated with or without ATP (500 μM, 25 μM) (S5260; Selleck Biotechnology.) for 30 min, followed by treatment with or without recombinant C5a (10 nM) (300-70-20UG; Thermo Fisher Scientific, Inc.) for 1 hour at 37 °C. Cells were then incubated with Sytox Green according to the manufacturer’s protocol (S7020; Thermo Fisher Scientific, Inc.). Fluorescence images were acquired using a confocal laser scanning microscope (FLUOVIEW FV10i; Olympus, Tokyo, Japan). Total neutrophils and dead cells were visualized in bright-field and Sytox Green staining, respectively. Sytox Green-positive areas were quantified using ImageJ with thresholding value of >127 (on a scale of 0–255). NETotic neutrophils were defined as cells with Sytox Green-stained areas > 68 μm², based on a previously established method [22]. NET frequency ((NETotic cells/total neutrophils) × 100) was calculated from the obtained images.

### 2.7. Scratch assay

HaCaT cells were seeded in 3.5 cm dish at 4.0 × 10^5^ cells/mL in DMEM. Following an incubation for 4 hours, cell monolayers were uniformly scratched using 1 mL microtips at 5 points along the diameter of the dish. Scratched HaCaT cells were incubated for 1 hour, and the concentrations of extracellular ATP, uridine triphosphate (UTP) and uridine diphosphate (UDP) in the supernatant were measured using Luminescent ATP Detection Assay Kit (ab113849; Abcam), ELISA Kit for Uridine Triphosphate (CEG822Ge: Cloud-Clone CORP., Katy, Texas, U.S.), and MicroMolar UDP assay kit (MUD100K; ProFoldin., Hudson, Massachusetts, U.S.) according to the manufacturer’s protocol.

### 2.8. Statistics

All experiments were analyzed using R (R Foundation for Statistical Computing, Vienna, Austria). Statistical significance was determined by Mann-Whitney U test or analysis of variance (ANOVA), and indicated by a bar and asterisk above each data set. A value of P <0.05 was considered significant. Where no significance was detected, no bar or asterisk is included in the figures.

## 3. Results

### 3.1. NETs inhibition decreases disease severity in EBA murine model

To assess the contribution of NETs in neutrophilic skin diseases, we utilized the passive IgG-transfer EBA murine model, administering intraperitoneal injections of rabbit anti-mouse Col7 IgG every two days for 12 days (Fig. 1A). Consistent with previous reports [20], mice developed skin lesions predominantly affecting the ears, head, face, neck, and legs, areas frequently scratched or bitten, accompanied by histological subepidermal blistering by day 12 (Fig. 1A-C). With immunohistochemistry, infiltrating cells were primarily neutrophils and NET formation was observed, as indicated by co-distribution of the neutrophilic enzyme, MPO, and CitH (Fig. 1D).

**Figure. 1.**
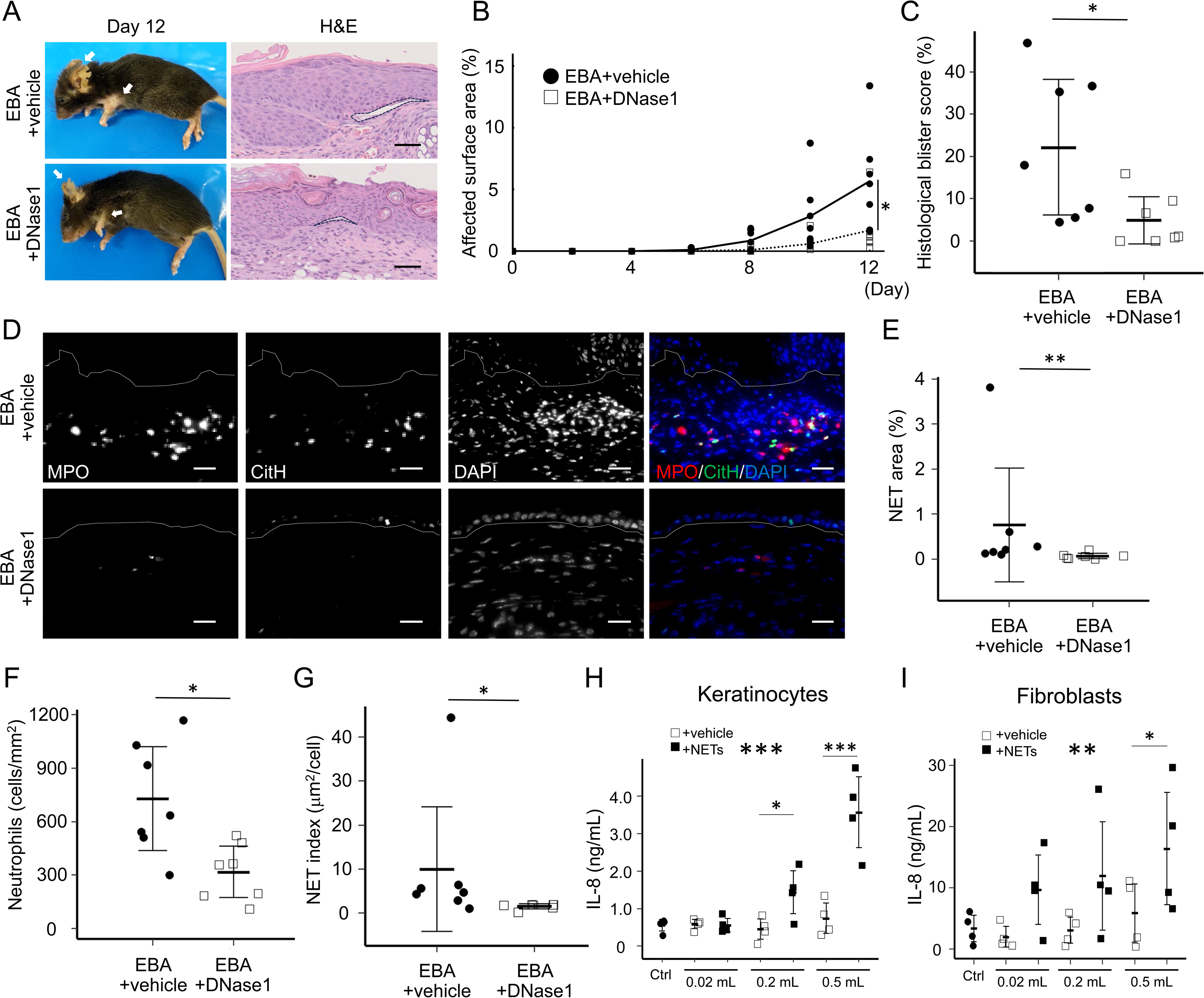
NETosis contributes to blistering in epidermolysis bullosa acquisita (EBA) murine model and induces IL-8 secretion from keratinocytes and fibroblasts in vitro. (A) Representative clinical images and H&E-stained ear sections from EBA mice treated with vehicle (EBA+vehicle) or DNase1 (EBA+DNase1) on day 12. White arrows indicate lesional skin. Dotted lines demarcate major dermal-epidermal separations. Scale bar = 100 µm. (B) Affected surface area defined as ((total affected body surface area)/(total body surface area) × 100) was quantified every 2 days for 12 days. Dot plots indicate all individual scores and lines indicate group means. N = 7 for each group. *P<0.05 (Two-way ANOVA (time and group as two variables)). (C) Histological blister scores defined as ((combined total length of all blistered regions)/(combined total length of all dermal-epidermal junction examined) × 100) were quantified from H&E staining images at day 12. Data shown as dot plots with mean ± standard deviation. N = 7 for each group. *P<0.05 (Mann-Whitney U test). (D) Representative immunohistochemistry of ear sections from EBA+vehicle and EBA+DNase1 mice stained for myeloperoxidase (MPO), citrullinated Histone H3 (CitH), and 4, 6-diamidino-2-phenylindole (DAPI). Co-distributed area of MPO and CitH indicates NETosis. Dotted lines indicate the dermal-epidermal junction. Scale bar = 40 µm. Quantification of NETosis area (E: (NETosis area of upper dermis)/(total area of upper dermis) × 100), infiltrating neutrophils (F: (neutrophil number in upper dermis)/(total area of upper dermis)), and NET index (G: (NETosis area of upper dermis)/(neutrophil number in upper dermis)). Data shown as dot plots with mean ± standard deviation. N = 7 for each group. *P<0.05, **P<0.01 (Mann-Whitney U test). IL-8 secretion by HaCaT cells (H) or fibroblasts (I) following 12-hour stimulation with 0.02, 0.2, or 0.5 mL of supernatant with or without NET components. IL-8 levels measured by ELISA. Data shown as dot plots with mean ± standard deviation. N = 4 for each group. *P<0.05, **P<0.01, ***P<0.001 (Two-way ANOVA (dose and group as two variables) with multiple comparison test). Large asterisks in the center indicate ANOVA significance, and small asterisks show pairwise comparisons.

To evaluate the pathological role of NETs in EBA, EBA mice were treated with DNase1 to degrade NET components. While DNase1-treated groups also developed skin lesions by day 12, the area of affected skin was significantly smaller than in vehicle controls, with mean affected body surface areas of 5.7% and 1.7% in the vehicle- and DNase1-treated groups, respectively (Fig. 1A, B). Similarly, histopathological analysis revealed reduced subepidermal blistering in DNase1-treated mice compared to the vehicle-treated group, with mean blister scores of 22.2% and 4.9%, respectively (Fig. 1A, C). With immunohistochemistry, DNase1 treatment was confirmed to reduce NETosis, as demonstrated by both NET area (from 0.76 to 0.07%) and NET index (NET area per infiltrating neutrophil numbers; from 10.0 to 1.5 μm²/cell) (Fig. 1D, E, G). Furthermore, neutrophilic infiltration was significantly decreased, with mean neutrophil numbers of 730 cells/ mm² and 318 cells/ mm² in vehicle- and DNase1-treated groups, respectively (fig. 1D, F). These findings support a pathological role of NETosis in EBA, potentially through the promotion of neutrophil recruitment.

### 3.2. NET components induce IL-8 secretion from keratinocytes and fibroblasts in vitro

To elucidate the mechanism by which NETs contribute to neutrophil recruitment in the EBA murine model, we assessed whether keratinocytes and fibroblasts stimulated with NET components release IL-8, a prominent neutrophil chemokine. To investigate this, cell-and cytokine-free NET components were isolated from PMA stimulated healthy neutrophils and were applied to HaCaT or fibroblasts. Both cell types exposed to purified NET components secreted significantly higher protein levels of IL-8 detected by ELISA in a dose-dependent manner (Fig. 1H, I). These results suggest that NET components can directly activate keratinocytes and fibroblasts, enhancing the production of IL-8, which contributes to further neutrophil recruitment. These findings suggest that NETs exhibit a pivotal role in EBA pathogenesis through IL-8 release from keratinocytes and fibroblasts resulting in further recruitment of neutrophils.

### 3.3. Mechanical stress reduction decreases disease severity and NETosis in EBA murine model

Since exaggerated inflammatory responses to minor mechanical stimulation are a hallmark of neutrophilic skin diseases and have also been reported in the EBA murine model [1, 3, 4, 23], we next tested whether mechanical stress contributes to NETosis and disease progression in this setting.

To address this, mice were applied neck-collars beginning 3 days prior to pathogenic IgG injection to restrict scratching. This intervention effectively decreased scratching of the facial skin, although it did not fully prevent mechanical stimulation of the neck and extremities due to collar friction, itch-related biting behavior, and movement-associated mechanical stress. Therefore, subsequent analysis focused on the facial skin. By day 12, both control and neck-collared EBA mice developed facial lesions, however, the lesional area was significantly smaller in the neck-collared mice, with mean facial lesional areas of 0.65% and 0.35%, respectively (Fig. 2A, B). Histological analysis of the ears confirmed reduced subepidermal blistering in neck-collared mice compared to the control, with mean blister scores of 37.8% and 3.4%, respectively (Fig. 2A, C). Immunohistochemical analysis further revealed that neck-collar treatment reduced the NET area (3.2% vs. 0.1%) and neutrophilic infiltration (719 cells/mm² vs. 205 cells/mm²) (Fig. 2D-F). Importantly, the NET index was also decreased in neck-collared mice (15.3 μm²/cell vs. 8.0 μm²/cell) (Fig. 2G). This indicates that the suppression of NET formation was not solely attributed by the reduced neutrophilic infiltration but also reflected lower susceptibility of NETosis. Collectively, these findings suggest that mechanical stress enhances NET formation and subsequent neutrophilic infiltration, thereby increasing disease severity in the EBA model.

**Figure. 2.**
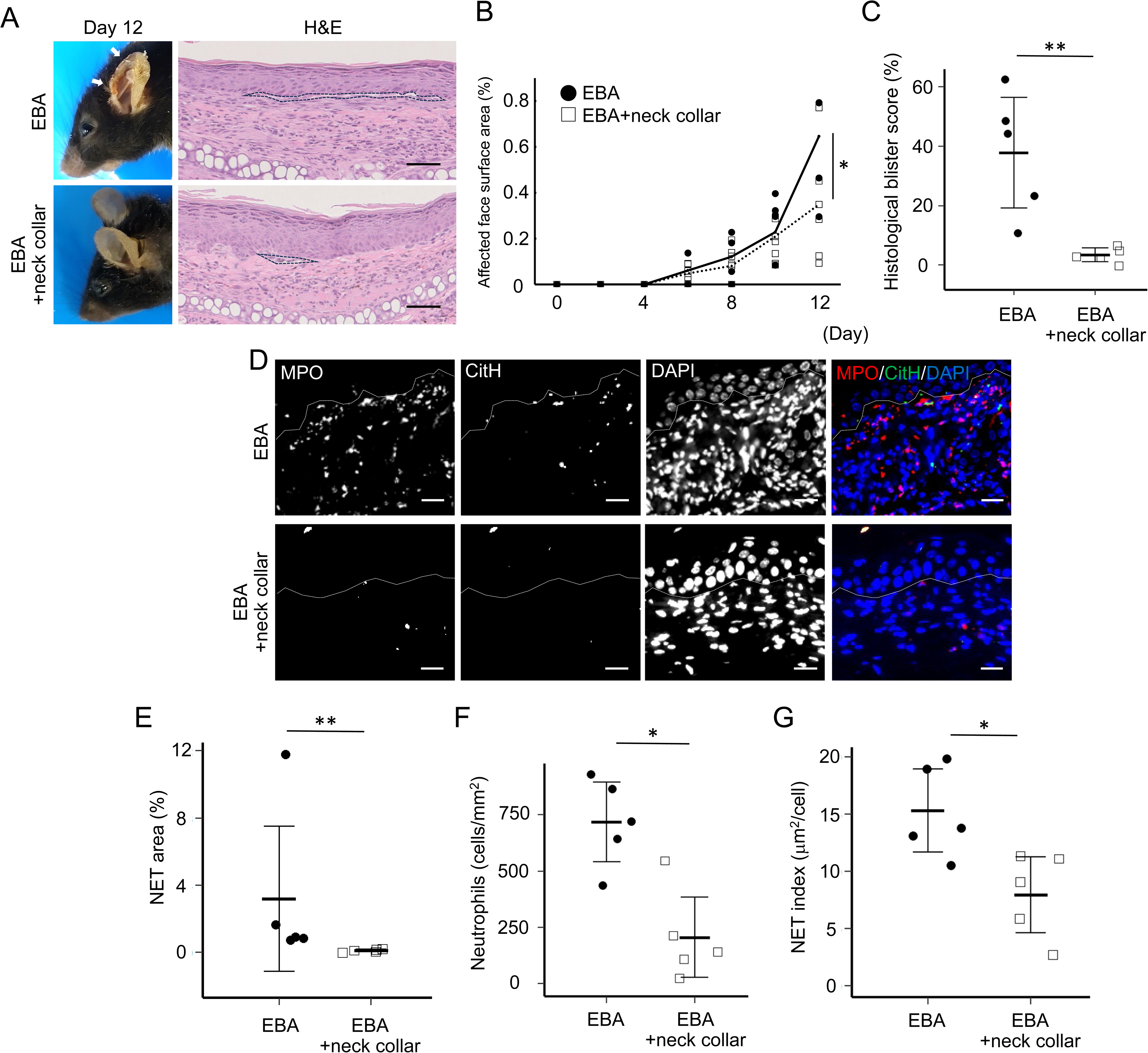
Restriction of scratching behavior by neck collar reduces blistering, neutrophilic infiltration, and NETosis in epidermolysis bullosa acquisita (EBA) murine model. (A) Representative clinical images and H&E-stained ear sections from EBA mice with or without neck collars (EBA and EBA+neck collar) on day 12. White arrows indicate lesional skin. Dotted lines demarcate major dermal–epidermal separations. Scale bar = 100 µm. (B) Affected face surface areas defined as ((total affected face surface area)/(total body surface area) × 100) were quantified every 2 days for 12 days. Dot plots indicate all individual scores and lines indicate group means. N=5 for each group. *P<0.05 (Two-way ANOVA (time and group as two variables)). (C) Histological blister scores defined as ((combined total length of all blistered regions)/(combined total length of all dermal-epidermal junction examined) × 100) were quantified from H&E staining images at day 12. Data shown as dot plots with mean ± standard deviation. N = 5 for each group. **P<0.01 (Mann-Whitney U test). (D) Representative immunohistochemistry of ear sections from EBA and EBA+neck collar mice stained for myeloperoxidase (MPO), citrullinated Histone H3 (CitH), and 4, 6-diamidino-2-phenylindole (DAPI). Co-distributed area of MPO and CitH indicates NETosis. Dotted lines indicate the dermal-epidermal junction. Scale bar = 40 µm. Quantification of NETosis area (E: (NETosis area of upper dermis)/(total area of upper dermis) × 100), infiltrating neutrophils (F: (neutrophil number in upper dermis)/(total area of upper dermis)), and NET index (G: (NETosis area of upper dermis)/(neutrophil number in upper dermis)). Data shown as dot plots with mean ± standard deviation. N = 5 for each group. *P<0.05, **P<0.01 (Mann-Whitney U test).

### 3.4. Inhibition of purinergic signaling decreases NET formation and disease severity in EBA murine model

Since keratinocytes constitute the outermost layer of the body, they are mechanosensitive and act as early transducers of external mechanical stimulations [24]. Based on previous reports showing that mechanical stress induces the release of extracellular nucleotides, including ATP, from keratinocytes [25–27], we examined whether keratinocyte-derived nucleotides released upon mechanical stimulation contribute to NETosis. Using an in vitro scratch assay, we confirmed that mechanical stress significantly increased ATP release from the keratinocytes (2.5 nM vs. 22.4 nM), whereas UTP and UDP levels remained unchanged (5.47 nM vs. 5.94 nM and 0.92 nM vs. 1.03 nM, respectively) (Fig. 3A-C).

**Figure. 3.**
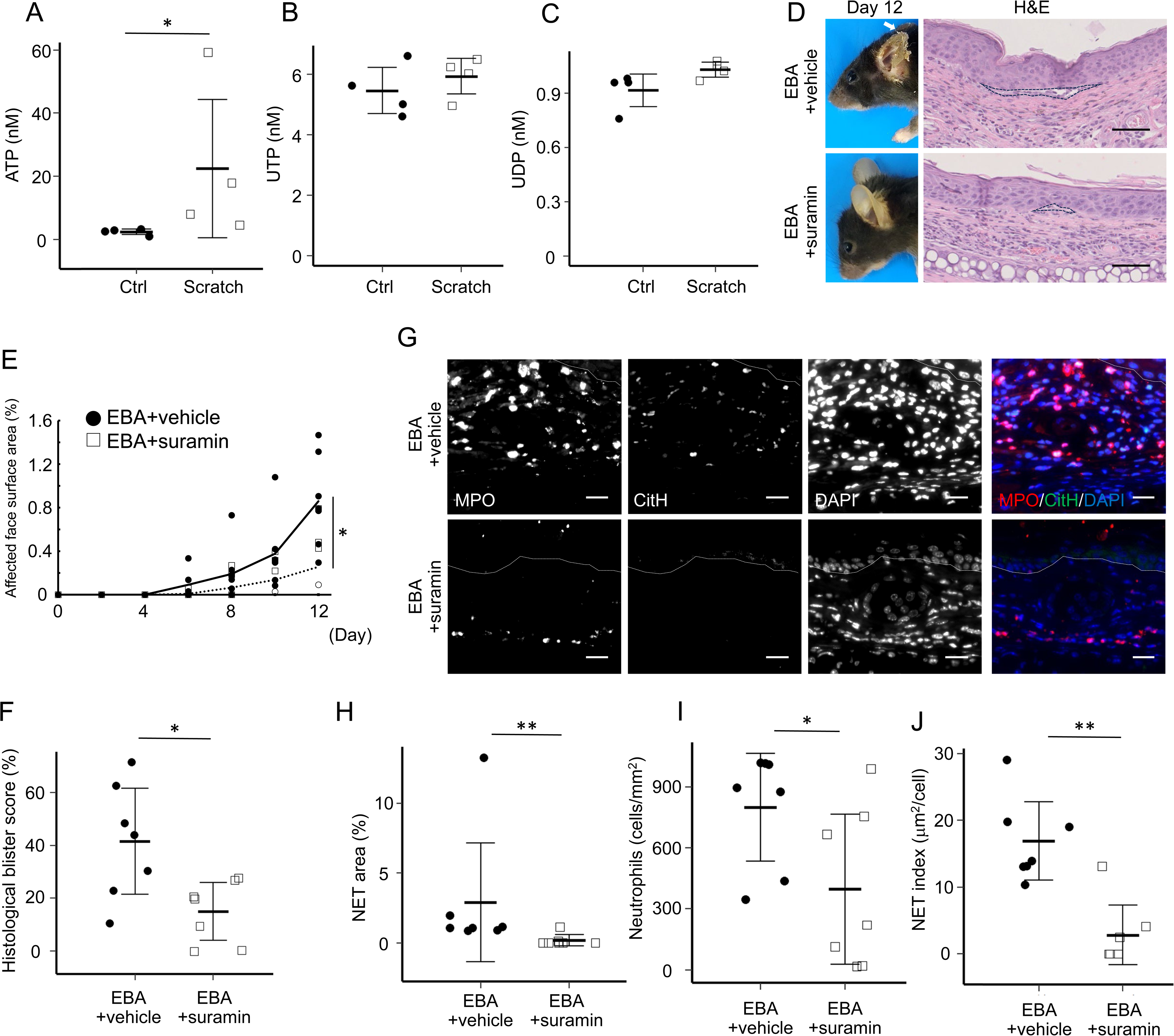
Purinergic signaling blockade reduces blistering, neutrophilic infiltration, and NETosis in epidermolysis bullosa acquisita (EBA) murine model. (A, B, C) ATP, UTP, and UDP in the supernatants of scratched keratinocytes were quantified by ELISA or bioluminescence assay. Data shown as dot plots with mean ± standard deviation. N = 4 for each group. *P<0.05 (Mann-Whitney U test). (D) Representative clinical images and H&E-stained ear sections from EBA mice treated with vehicle (EBA+vehicle) or suramin (EBA+suramin) on day 12. White arrow indicates lesional skin. Dotted lines demarcate major dermal–epidermal separations. Scale bar = 100 µm. (E) Affected face surface area defined as ((total affected face surface area)/(total body surface area) × 100) was quantified every 2 days for 12 days. Dot plots indicate all individual scores and lines indicate group means. N = 7 for each group. *P<0.05 (Two-way ANOVA (time and group as two variables)). (F) Histological blister scores defined as ((combined total length of all blistered regions)/(combined total length of all dermal-epidermal junction examined) × 100) were quantified from H&E staining images at day 12. Data shown as dot plots with mean ± standard deviation. N = 7 for each group. *P<0.05 (Mann-Whitney U test). (G) Representative immunohistochemistry of ear sections from EBA+vehicle and EBA+suramin mice stained for myeloperoxidase (MPO), citrullinated Histone H3 (CitH), and 4ʹ,6-diamidino-2-phenylindole (DAPI). Co-distributed area of MPO and CitH indicates NETosis. Dotted lines indicate the dermal-epidermal junction. Scale bar = 40 µm. Quantification of NETosis area (H: (NETosis area of upper dermis)/(total area of upper dermis) × 100), infiltrating neutrophils (I: (neutrophil number in upper dermis)/(total area of upper dermis)), and NET index (J: (NETosis area of upper dermis)/(neutrophil number in upper dermis)). Data shown as dot plots with mean ± standard deviation. N = 7 for each group. *P<0.05, **P<0.01 (Mann-Whitney U test).

To assess whether released ATP following mechanical stress contributes to NET formation to increase disease severity in EBA, we administered suramin, a purinergic receptor antagonist, into the EBA mice intraperitoneally for 12 consecutive days, starting one day before anti-Col7 IgG administration. By day 12, while both vehicle- and suramin-treated groups developed lesions, the affected facial area was significantly smaller in suramin-treated mice, with mean facial areas affected of 0.86% and 0.26%, respectively (Fig. 3D, E). As observed with the neck-collar intervention, suramin treatment did not decrease lesions on the trunk and extremities, likely due to continued itch-related biting behavior and movement-associated mechanical stimulations. Histological analysis of the ear tissue further confirmed reduced subepidermal blistering in suramin-treated mice, with mean histological blistering scores of 41.6% and 15.0% in vehicle- and suramin-treated mice, respectively (Fig. 3D, F). In addition, immunohistochemical analyses revealed a significant decrease in NET area, neutrophilic infiltration, and NET index of suramin-treated mice compared to controls, with NET area of 3.0% vs. 0.19%, neutrophil densities of 800 vs. 397 cells/mm², and NET index of 17.0 μm²/cell vs. 2.8 μm²/cell, respectively (Fig. 3H-J). These findings suggest that the inhibition of signaling from extracellular nucleotide signaling suppresses NETosis and subsequent neutrophil recruitment, leading to reduced blister formation.

### 3.5. ATP enhances C5a-induced NETosis in vitro

Based on these observations, extracellular nucleotides, particularly ATP, were suggested to contribute to NETosis. To test their direct contribution, healthy human neutrophils were stimulated with ATP in vitro (Fig. 4A, B). However, ATP alone did not induce significant NETosis, defined as cells showing Sytox Green stained areas larger than 68 μm², a threshold previously reported to distinguish NETotic from apoptotic cells [22]. This result demonstrated that ATP alone does not trigger NETosis, consistent with previous findings [28].

**Figure. 4.**
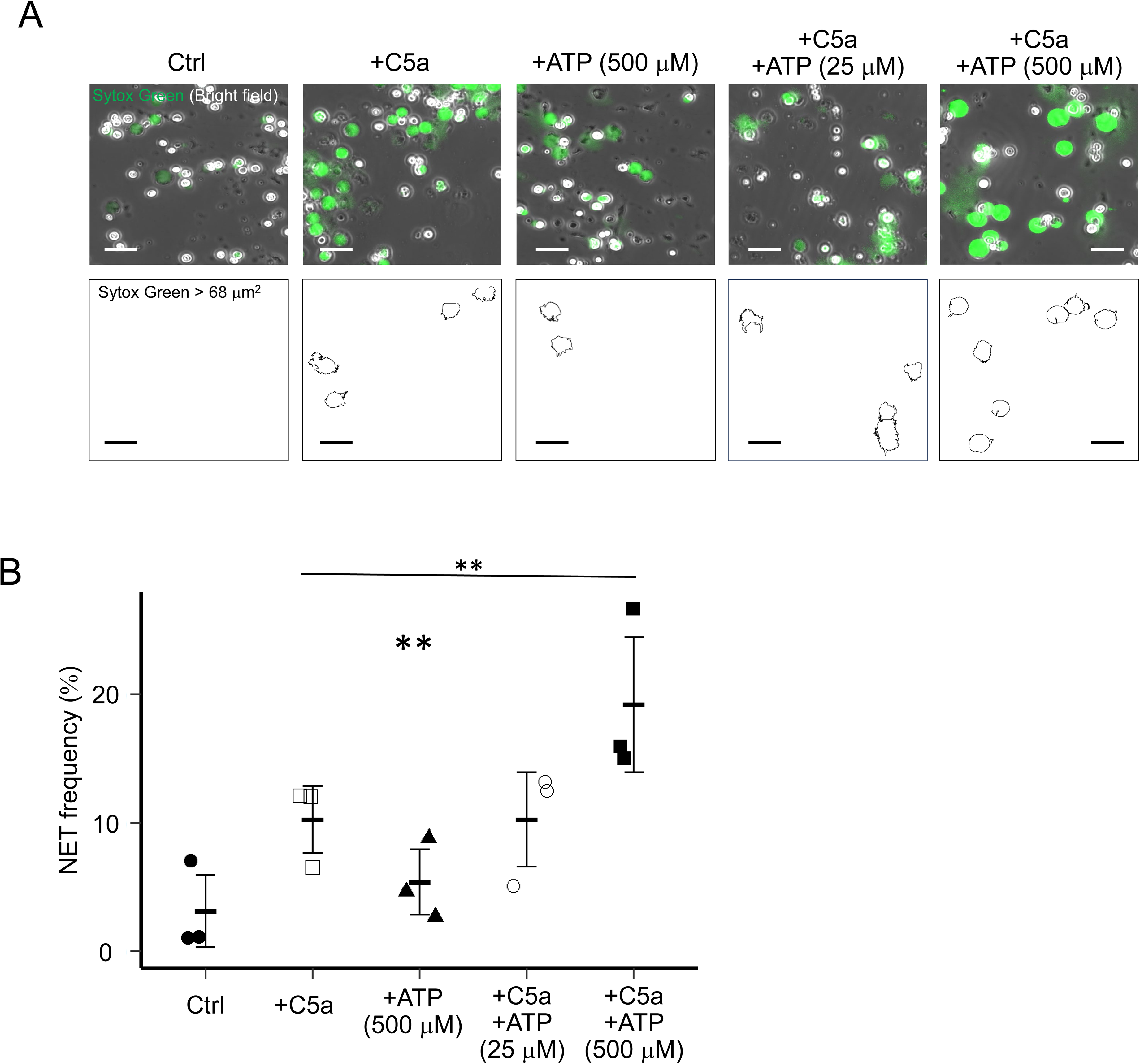
ATP stimulation on neutrophils enhances NETosis in the presence of activated complements. (A) Upper panels show representative merged images of Sytox Green fluorescence and bright-field microscopy of neutrophils pre-treated with or without ATP, followed by stimulation with or without recombinant C5a. Lower panels display binary images highlighting Sytox Green-positive areas greater than 68 µm2, defined as NETotic cells. Scale bar = 40 µm. (B) Quantification of NET frequencies defined as ((NETotic cells/neutrophils) × 100). Data shown as dot plots with mean ± standard deviation. N = 3 for each group. **P<0.01 (pairwise multiple comparison test following one-way ANOVA). Large asterisks in the center indicate ANOVA significance, and small asterisks show pairwise comparisons.

Since direct stimulation was insufficient, we next evaluated whether ATP stimulation indirectly promotes NET formation within inflammatory microenvironment. Because the anaphylatoxin C5a signaling drives EBA blistering and NETosis [29], we tested the involvement of ATP in combination with C5a. Healthy human neutrophils were stimulated with human recombinant C5a, with or without the pretreatment with ATP (Fig. 4A, B). While minimal NETosis was observed with C5a alone, ATP supplementation markedly increased NET formation (10.2% vs. 19.2%).

These results suggest that extracellular ATP enhances C5a-induced NET formation. In the context of EBA, ATP release from mechanically stressed keratinocytes may enhance NETosis by amplifying the effects of inflammatory cues such as C5a in the lesional skin.

### 3.6. Prominent NET formation in the upper dermis of neutrophilic skin disorders

Finally, we evaluated the distribution of NETs in various neutrophilic skin diseases, including EBA (N = 6), Behçet disease (N = 3), Sweet syndrome (N = 5), and PG (N = 5), compared with healthy controls (N = 3). In all four diseases, NETs were detected among infiltrating neutrophils, although the frequency of NET formation varied between diseases (Fig. 5A, B). Notably, NETs were more abundant in the upper dermis than in the lower dermis in EBA and PG, mirroring findings from the EBA murine model. These findings indicate that NETosis is induced within skin-infiltrating neutrophils in neutrophilic dermatoses and that NETs are preferentially induced at the upper dermis in EBA and PG. These results support a model in which keratinocyte-derived ATP contribute to localized NETosis in the upper dermis, thereby promoting further neutrophilic infiltration and amplifying inflammation in these neutrophilic skin diseases, particularly in EBA and PG.

**Figure. 5.**
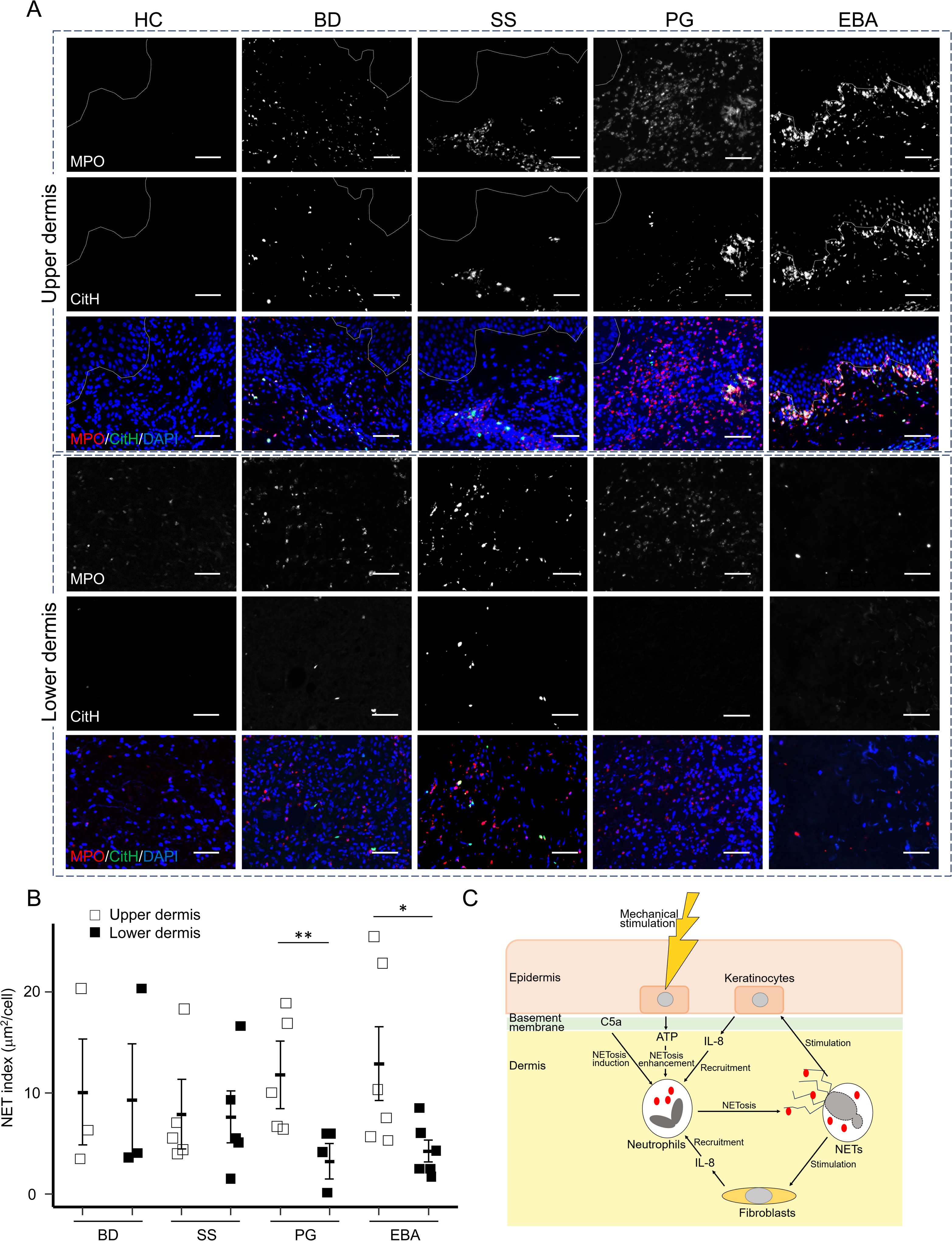
Prominent NET formation in human neutrophilic skin disorders. (A) Representative immunohistochemistry of human skin sections from healthy control (HC), Behçet disease (BD), Sweet syndrome (SS), pyoderma gangrenosum (PG), and epidermolysis bullosa acquisita (EBA) for myeloperoxidase (MPO), citrullinated Histone H3 (CitH), and 4ʹ,6-diamidino-2-phenylindole (DAPI). Dotted lines indicate the dermal-epidermal junction. Scale bar = 40 µm. (B) Quantification of NET index (NETosis area of upper or lower dermis)/(Neutrophil number in upper or lower dermis). The dot plots indicate all individual scores and error bars indicate mean ± standard deviation. N = 3 (HC), 3 (BD), 5 (SS), 5 (PG), and 6 (EBA). *P<0.05, **P<0.01 (Mann-Whitney U test). (C) Proposed model of the mechanical stimulation-ATP-NETs axis in EBA. Mechanical stimulation triggers ATP release from keratinocytes. In combination with C5a, extracellular ATP promotes NETosis. NET components induce IL-8 release from keratinocytes and fibroblasts, resulting in further neutrophil recruitment.

## 4. Discussion

Neutrophilic skin diseases, including Behçet disease, Sweet syndrome, PG, and EBA, exhibit a common clinical feature: exaggerated inflammatory responses to minor physical stimulation on the skin. While phenomena such as positive pathergy test in Behçet disease, trauma induced lesion formation in PG and Sweet syndrome, and friction-induced blistering in EBA are well documented, the underlying mechanisms remain poorly understood.

In this manuscript, we elucidated a mechanistic link between mechanical skin stimulation and exaggerated neutrophilic inflammation in EBA, mediated by keratinocyte-derived ATP and NETosis. Specifically, we demonstrated that (1) NETosis contributes to disease severity in EBA, (2) mechanical stress induces ATP release from keratinocytes, which in combination with activated complement, promotes NETosis, (3) NET components create a self-amplifying loop that sustains neutrophilic infiltration via IL-8 release from keratinocytes and fibroblasts, and (4) NETosis localizes to the upper dermis in EBA and PG.

Several alternative mechanisms by which mechanical stress exacerbates autoantibody-mediated skin diseases have been already proposed. For example, skin friction has been revealed to increase vascular permeability, facilitating pathogenic IgG access to the dermal-epidermal junction in pemphigoid disease model [30]. On the other hand, scratching has been indicated to alter the cytokine milieu, skewing inflammation toward Th1 polarization in EBA model [23]. However, these studies do not explain the predominance of hypersensitivity observed in neutrophilic skin diseases. Our findings provide a distinct perspective by identifying NETosis as a central driver of this hypersensitivity, independent from IgG-mediated mechanisms.

Previous studies indicate that mechanical stimulations to the skin are sensed primarily by keratinocytes and transduced through ATP release [25–27]. For example, previous studies have shown that stretched keratinocytes release ATP in vitro, which in turn activates immune cells and keratinocytes [25]. This mechanical-stretch model utilized by Okamoto and his colleagues may better reflect in vivo conditions than traditional in vitro scratch assays, especially in the context of in vivo scratching in pruritic skin diseases. These findings align with our results, supporting a model in which epidermal mechanical stimulation, such as scratch or friction, leads to ATP release from keratinocytes to promote NETosis.

Mechanistically, we demonstrate that ATP alone does not trigger NETosis, but potentiates C5a-mediated NETosis in vitro. Conversely, low concentrations of C5a only induce minimal NETosis, while co-stimulation with ATP nearly doubled NETosis. In vivo, we further demonstrated that inhibition of ATP signaling decreases both lesional NETosis and blistering in the EBA model, indicating that ATP signaling is required for full disease expression under physiological levels of activated complements. Together, these data support that ATP signaling acts as a second hit to amplify an ongoing adaptive to innate immune cascade. These data support a model in which complement activation primes neutrophils, while tissue-derived stress signals provide the necessary context for NET release. While former studies have already shown that NETosis serves as a mechanistic bridge between humoral and innate immunity via complement activation [9], our data highlight how lesional NETosis following humoral immunity is regulated tightly. Moreover, once NETs are formed, NETs themselves exaggerate inflammation by inducing IL-8 release from keratinocytes and fibroblasts, creating a self-reinforcing loop of neutrophil recruitment and activation.

We also observed a consistent pattern of NET localization in skin samples from EBA and PG, where NETs are predominantly localized in the upper dermis. This finding suggests that shared triggering factors, ATP released from nearby keratinocytes (rather than fibroblasts) and activated complement are involved in inducing NETosis in both diseases. This is consistent with the former reports identifying both EBA and PG as complement-associated neutrophilic skin disease [31, 32]. Since psoriasis has also been reported to involve complement activation and NETosis [33, 8], similar ATP- and complement-mediated mechanisms may contribute to its pathology and to its characteristic hypersensitivity to the skin stimulation, known as Koebner phenomenon. In contrast, deeper dermal NETosis was observed in other neutrophilic skin diseases such as Behçet disease and Sweet syndrome. These deeper NETs may reflect mechanisms independent of both complement and keratinocyte-released ATP. Further investigation is needed to elucidate these pathways and to define the cellular sources responsible to site-specific NET induction in these neutrophilic skin diseases. Beyond neutrophilic diseases, mechanical stress by scratching is a well-established aggravating factor in other pemphigoid diseases, particularly in bullous pemphigoid (BP) [34]. This suggests that the ATP–extracellular trap formation axis may contribute more broadly to the pathogenesis of pemphigoid diseases. Notably, while neutrophilic infiltration dominates in EBA, BP is characterized by prominent eosinophilic infiltration [35], suggesting that eosinophil extracellular trap formation (EETosis), as previously observed [36], may be involved.

From a therapeutic perspective, our data suggest that targeting components of the ATP-NET axis may attenuate disease activity in neutrophilic skin diseases including EBA. Furthermore, the link between mechanical stress and inflammation underscores the importance of controlling itch. Indeed, our findings provide new insight into the mechanism underlying itch-scratch cycle in neutrophilic pruritic skin diseases.

In conclusion, this study establishes a mechanistic framework in which keratinocyte-derived ATP links mechanical stress to NETosis, thereby integrating signals from humoral and innate immunity. This axis not only drives chronic inflammation in EBA but may also operate in a broader spectrum of neutrophilic skin disorders including PG. By identifying this self-amplifying cascade, we highlight novel targets for interrupting disease progression in these challenging conditions.

## Study approval

The privacy rights of the patients were respected, and informed consent for the use of clinical test data was obtained in accordance with the principles of the Declaration of Helsinki.

## Acknowledgments

We express our thanks to Mr. Keisuke Inoue and Ms. Emi Donoue from the Research Support Platform in Graduate School of Medicine at Osaka Metropolitan University for their assistance with sample preparation for immunohistochemistry.

## CRediT authorship contribution statement

**Daisuke Matsumoto:** Formal analysis, Investigation, Writing – original draft & review & editing. **Katja Bieber:** Funding acquisition, Methodology, Resources, Writing – review & editing. **Ralf J. Ludwig:** Funding acquisition, Methodology, Resources, Writing – review & editing. **Daisuke Tsuruta:** Funding acquisition, Writing – review & editing. **Sho Hiroyasu:** Conceptualization, Funding acquisition, Project administration, Supervision, Writing – original draft & review & editing.

## Declaration of generative AI and AI-assisted technologies in the writing process

During the preparation of this work, the authors used ChatGPT to improve the language in some individual sentences only. After using this tool/service, the author(s) reviewed and edited the content as needed and take(s) full responsibility for the content of the published article.

## Data availability

Data will be made available on request.

## References

[1] A. V. Marzano, A. Borghi, D. Wallach, M. Cugno, A Comprehensive Review of Neutrophilic Diseases, Clin Rev Allergy Immunol 54 (2018) 114–130. 10.1007/s12016-017-8621-8.

[2] K. Kridin, D. Kneiber, E.H. Kowalski, M. Valdebran, K.T. Amber, Epidermolysis bullosa acquisita: A comprehensive review, Autoimmun Rev 18 (2019) 786–795. 10.1016/j.autrev.2019.06.007.

[3] C.A. Nelson, S. Stephen, H.J. Ashchyan, W.D. James, R.G. Micheletti, M. Rosenbach, Neutrophilic dermatoses: Pathogenesis, Sweet syndrome, neutrophilic eccrine hidradenitis, and Behçet disease, J Am Acad Dermatol 79 (2018) 987–1006. 10.1016/j.jaad.2017.11.064.

[4] A. Varol, O. Seifert, C.D. Anderson, The skin pathergy test: Innately useful?, Arch Dermatol Res 302 (2010) 155–168. 10.1007/s00403-009-1008-9.

[5] S. Li, S. Ying, Y. Wang, Y. Lv, J. Qiao, H. Fang, Neutrophil extracellular traps and neutrophilic dermatosis: an update review, Cell Death Discov 10 (2024). 10.1038/s41420-023-01787-2.

[6] V. Brinkmann, U. Reichard, C. Goosmann, B. Fauler, Y. Uhlemann, D.S. Weiss, Y. Weinrauch, A. Zychlinsky, Neutrophil Extracellular Traps Kill Bacteria, Science 303 (2004) 1532–1535. 10.1126/science.1092385.

[7] Q. Remijsen, T. Vanden Berghe, E. Wirawan, B. Asselbergh, E. Parthoens, R. De Rycke, S. Noppen, M. Delforge, J. Willems, P. Vandenabeele, Neutrophil extracellular trap cell death requires both autophagy and superoxide generation, Cell Res 21 (2011) 290–304. 10.1038/cr.2010.150.

[8] F. Herster, Z. Bittner, N.K. Archer, S. Dickhöfer, D. Eisel, T. Eigenbrod, T. Knorpp, N. Schneiderhan-Marra, M.W. Löffler, H. Kalbacher, T. Vierbuchen, H. Heine, L.S. Miller, D. Hartl, L. Freund, K. Schäkel, M. Heister, K. Ghoreschi, A.N.R. Weber, Neutrophil extracellular trap-associated RNA and LL37 enable self-amplifying inflammation in psoriasis, Nat Commun 11 (2020). 10.1038/s41467-019-13756-4.

[9] M. Maqsood, S. Suntharalingham, M. Khan, C.G. Ortiz-Sandoval, W.J.C. Feitz, N. Palaniyar, C. Licht, Complement-Mediated Two-Step NETosis: Serum-Induced Complement Activation and Calcium Influx Generate NADPH Oxidase-Dependent NETs in Serum-Free Conditions, Int J Mol Sci 25 (2024) 9625. 10.3390/ijms25179625.

[10] Y. Wang, M. Li, S. Stadler, S. Correll, P. Li, D. Wang, R. Hayama, L. Leonelli, H. Han, S.A. Grigoryev, C.D. Allis, S.A. Coonrod, Histone hypercitrullination mediates chromatin decondensation and neutrophil extracellular trap formation, J Cell Biol 184 (2009) 205–213. 10.1083/jcb.200806072.

[11] E. Neubert, D. Meyer, F. Rocca, G. Günay, A. Kwaczala-Tessmann, J. Grandke, S. Senger-Sander, C. Geisler, A. Egner, M.P. Schön, L. Erpenbeck, S. Kruss, Chromatin swelling drives neutrophil extracellular trap release, Nat Commun 9 (2018). 10.1038/s41467-018-06263-5.

[12] C.K. Smith, M.J. Kaplan, The role of neutrophils in the pathogenesis of systemic lupus erythematosus, Curr Opin Rheumatol 27 (2015) 448–453. 10.1097/BOR.0000000000000197.

[13] R. Khandpur, C. Carmona-Rivera, A. Vivekanandan-Giri, A. Gizinski, S. Yalavarthi, J.S. Knight, S. Friday, S. Li, R.M. Patel, V. Subramanian, P. Thompson, P. Chen, D.A. Fox, S. Pennathur, M.J. Kaplan, NETs are a source of citrullinated autoantigens and stimulate inflammatory responses in rheumatoid arthritis, Sci Transl Med 5 (2013). 10.1126/scitranslmed.3005580.

[14] M. Kinoshita, Y. Ogawa, N. Hama, I. Ujiie, A. Hasegawa, S. Nakajima, T. Nomura, J. Adachi, T. Sato, S. Koizumi, S. Shimada, Y. Fujita, H. Takahashi, Y. Mizukawa, T. Tomonaga, K. Nagao, R. Abe, T. Kawamura, Neutrophils initiate and exacerbate Stevens-Johnson syndrome and toxic epidermal necrolysis, Sci Transl Med 13 (2021). 10.1126/scitranslmed.aax2398.

[15] H. Fang, S. Shao, K. Xue, X. Yuan, P. Qiao, J. Zhang, T. Cao, Y. Luo, X. Bai, W. Li, C. Li, H. Qiao, E. Dang, G. Wang, Neutrophil extracellular traps contribute to immune dysregulation in bullous pemphigoid via inducing B-cell differentiation and antibody production, FASEB J 35 (2021). 10.1096/fj.202100145R.

[16] D. Matsumoto, B. Amatya, D. Tsuruta, S. Hiroyasu, The pathological function of neutrophils in pemphigoid diseases, Dermatol Sin 42 (2024) 80–88. 10.4103/ds.DS-D-24-00027.

[17] K. Bieber, R.J. Ludwig, A. Kasprick, Drug Discovery for Pemphigoid Diseases, Curr Protoc Pharmacol 84 (2019) 55. 10.1002/cpph.55.

[18] S. Mihai, M.T. Chiriac, K. Takahashi, J.M. Thurman, M. Holers, D. Zillikens, M. Botto, C. Sitaru, The Alternative Pathway of Complement Activation Is Critical for Blister Induction in Experimental Epidermolysis Bullosa Acquisita1, J Immunol 178 (2007) 6514–6521. 10.4049/jimmunol.178.10.6514.

[19] D. Giusti, E. Bini, C. Terryn, K. Didier, S. Le Jan, G. Gatouillat, A. Durlach, S. Nesmond, C. Muller, P. Bernard, F. Antonicelli, B.N. Pham, Net formation in bullous pemphigoid patients with relapse is modulated by IL-17 and IL-23 interplay, Front Immunol 10 (2019). 10.3389/fimmu.2019.00701.

[20] S. Hiroyasu, M.R. Zeglinski, H. Zhao, M.A. Pawluk, C.T. Turner, A. Kasprick, C. Tateishi, W. Nishie, A. Burleigh, P.A. Lennox, N. Van Laeken, N.J. Carr, F. Petersen, R.I. Crawford, H. Shimizu, D. Tsuruta, R.J. Ludwig, D.J. Granville, Granzyme B inhibition reduces disease severity in autoimmune blistering diseases, Nat Commun 12 (2021). 10.1038/s41467-020-20604-3.

[21] H. Kawasaki, K. Tsunoda, T. Hata, K. Ishii, T. Yamada, M. Amagai, Synergistic pathogenic effects of combined mouse monoclonal anti-desmoglein 3 IgG antibodies on pemphigus vulgaris blister formation, J Invest Dermatol 126 (2006) 2621–2630. 10.1038/sj.jid.5700450.

[22] M. Van Der Linden, G.H.A. Westerlaken, M. Van Der Vlist, J. Van Montfrans, L. Meyaard, Differential Signalling and Kinetics of Neutrophil Extracellular Trap Release Revealed by Quantitative Live Imaging, Sci Rep 7 (2017). 10.1038/s41598-017-06901-w.

[23] M. Niebuhr, K. Bieber, D. Banczyk, S. Maass, S. Klein, M. Becker, R. Ludwig, D. Zillikens, J. Westermann, K. Kalies, Epidermal Damage Induces Th1 Polarization and Defines the Site of Inflammation in Murine Epidermolysis Bullosa Acquisita, J Invest Dermatol 140 (2020) 1713–1722.e9. 10.1016/j.jid.2020.01.022.

[24] M. Denda, P.M. Elias, Review of sensory systems deployed by epidermal keratinocytes, Front Cell Dev Biol 13 (2025). 10.3389/fcell.2025.1598326.

[25] T. Okamoto, Y. Ogawa, M. Kinoshita, T. Ihara, S. Shimada, S. Koizumi, T. Kawamura, Mechanical stretch-induced ATP release from keratinocytes triggers Koebner phenomenon in psoriasis, J Dermatol Sci 103 (2021) 60–62. 10.1016/j.jdermsci.2021.06.001.

[26] F. Moehring, A.M. Cowie, A.D. Menzel, A.D. Weyer, M. Grzybowski, T. Arzua, A.M. Geurts, O. Palygin, C.L. Stucky, Keratinocytes mediate innocuous and noxious touch via ATP-P2X4 signaling, Elife 16 (2018). 10.7554/eLife.31684

[27] N. Azorin, M. Raoux, L. Rodat-Despoix, T. Merrot, P. Delmas, M. Crest, ATP signalling is crucial for the response of human keratinocytes to mechanical stimulation by hypo-osmotic shock, Exp Dermatol 20 (2011) 401–407. 10.1111/j.1600-0625.2010.01219.x.

[28] A. Sofoluwe, M. Bacchetta, M. Badaoui, B.R. Kwak, M. Chanson, ATP amplifies NADPH-dependent and -independent neutrophil extracellular trap formation, Sci Rep 9 (2019). 10.1038/s41598-019-53058-9.

[29] Y. Chen, X. Li, X. Lin, H. Liang, X. Liu, X. Zhang, Q. Zhang, F. Zhou, C. Yu, L. Lei, J. Xiu, Complement C5a induces the generation of neutrophil extracellular traps by inhibiting mitochondrial STAT3 to promote the development of arterial thrombosis, Thromb J 20 (2022). 10.1186/s12959-022-00384-0.

[30] J.E. Hundt, H. Iwata, M. Pieper, R. Pfündl, K. Bieber, D. Zillikens, P. König, R.J. Ludwig, Visualization of autoantibodies and neutrophils in vivo identifies novel checkpoints in autoantibody-induced tissue injury, Sci Rep 10 (2020). 10.1038/s41598-020-60233-w.

[31] S. Mihai, M. Hirose, Y. Wang, J.M. Thurman, V. Michael Holers, B. Paul Morgan, J. Köhl, D. Zillikens, R.J. Ludwig, F. Nimmerjahn, Specific inhibition of complement activation significantly ameliorates autoimmune blistering disease in mice, Front Immunol 9 (2018). 10.3389/fimmu.2018.00535.

[32] Z. Wang, N. Hornick, M. Vague, D. Yang, J. Keller, S. Kody, S. Leachman, A.G. Ortega-Loayza, Y. Liu, NETosis Is Induced by Complement Component 5a: Implications in the Pathogenesis of Pyoderma Gangrenosum, J Invest Dermatol 144 (2024) 184–188.e2. 10.1016/j.jid.2023.06.204.

[33] Q. Zheng, F. Xu, Y. Yang, D. Sun, Y. Zhong, S. Wu, G. Li, W. Gao, T. Wang, G. Xu, S. Liang, C5a/C5aR1 mediates IMQ-induced psoriasiform skin inflammation by promoting IL-17A production from γδ-T cells, FASEB Journal 34 (2020) 10590–10604. 10.1096/fj.202000384R.

[34] S. Hiroyasu, J.V.J.G. Barit, A. Hiroyasu, D. Tsuruta, Pruritogens in pemphigoid diseases: Possible therapeutic targets for a burdensome symptom, J Dermatol 50 (2023) 150–161. 10.1111/1346-8138.16652.

[35] K.T. Amber, M. Valdebran, K. Kridin, S.A. Grando, The role of eosinophils in bullous pemphigoid: A developing model of eosinophil pathogenicity in mucocutaneous disease, Front Med (Lausanne) 5 (2018). 10.3389/fmed.2018.00201.

[36] S. Shen, H. Fang, X. Li, Y. Zhou, X. Tang, H. Miao, L. Li, J. Chen, K. Xue, C. Zhang, M. Chu, B. Pang, Y. Bai, H. Qiao, E. Dang, S. Shao, G. Wang, Eosinophil extracellular traps drive T follicular helper cell differentiation via VIRMA-dependent MAF stabilization in bullous pemphigoid, J Allergy Clin Immunol 155 (2024) 1357–1370. 10.1016/j.jaci.2024.09.030.

